# Enhancing droplet-based single-nucleus RNA-seq resolution using the semi-supervised machine learning classifier DIEM

**DOI:** 10.1101/786285

**Authors:** Marcus Alvarez, Elior Rahmani, Brandon Jew, Kristina M. Garske, Zong Miao, Jihane N. Benhammou, Chun Jimmie Ye, Joseph R. Pisegna, Kirsi H. Pietiläinen, Eran Halperin, Päivi Pajukanta

## Abstract

Single-nucleus RNA sequencing (snRNA-seq) measures gene expression in individual nuclei instead of cells, allowing for unbiased cell type characterization in solid tissues. Contrary to single-cell RNA seq (scRNA-seq), we observe that snRNA-seq is commonly subject to contamination by high amounts of extranuclear background RNA, which can lead to identification of spurious cell types in downstream clustering analyses if overlooked. We present a novel approach to remove debris-contaminated droplets in snRNA-seq experiments, called Debris Identification using Expectation Maximization (DIEM). Our likelihood-based approach models the gene expression distribution of debris and cell types, which are estimated using EM. We evaluated DIEM using three snRNA-seq data sets: 1) human differentiating preadipocytes *in vitro*, 2) fresh mouse brain tissue, and 3) human frozen adipose tissue (AT) from six individuals. All three data sets showed various degrees of extranuclear RNA contamination. We observed that existing methods fail to account for contaminated droplets and led to spurious cell types. When compared to filtering using these state of the art methods, DIEM better removed droplets containing high levels of extranuclear RNA and led to higher quality clusters. Although DIEM was designed for snRNA-seq data, we also successfully applied DIEM to single-cell data. To conclude, our novel method DIEM removes debris-contaminated droplets from single-cell-based data fast and effectively, leading to cleaner downstream analysis. Our code is freely available for use at https://github.com/marcalva/diem.

## Introduction

Single-cell RNA sequencing (scRNA-seq) has grown considerably in use over the previous decade and permitted a transcriptomic view into the composition of heterogeneous mixtures of cells^1,2^. Recent advances in droplet-based microfluidics have created a high-throughput opportunity to assay single cells by scaling up previous well-based technologies to tens to hundreds of thousands of cells^3^. Single-nucleus RNA sequencing (snRNA-seq), where nuclei are used instead of cells, has allowed the critical extension of single-cell based technologies to solid tissues where isolation and suspension of individual cells is difficult or impossible^4^. For example, snRNA-seq has been used successfully to identify cell types in the brain^5^. Another practically important application of sequencing nuclei is identifying cell types in frozen tissue, from which it is often not feasible to isolate intact cells, whereas nuclei can still be successfully isolated^6^.

Droplet-based snRNA-seq encapsulates individual nuclei into a water droplet with reagents for generating cDNA and ligating droplet-specific oligonucleotide barcodes. After library construction and sequencing, the mapped reads can be assigned to droplets of origin. The input nuclei suspension is prepared so that all reads associated with one barcode originate from one nucleus. However, RNA originating from lysed cellular components (such as the cytoplasm) or from outside the cell can become encapsulated in droplets as well. Since these reads have the same barcode, contaminated RNA cannot be readily distinguished from nuclear RNA. To apply snRNA-seq to tissues, homogenization of the tissue is usually required to break apart the extracellular matrix and release nuclei from cells^4^. This can release higher amounts of debris and lead to more background RNA contamination^7^. This contamination of droplets with extranuclear RNA can lead to a biased increase in expression of these genes. Using mitochondrial RNA, we show that this results in clusters driven by background RNA, as well as contamination of clusters representing true cell types. As droplet-based snRNA-seq is increasingly applied to various solid tissues, there is an urgent need to accurately filter contaminated droplets.

A common practice to distinguish cell/nuclei-vs. background-containing barcodes relies on removing droplets below a hard cutoff of the number of reads, unique molecular indexes (UMI), or genes detected in a droplet^3,8–11^. This *ad hoc* cutoff is typically set by ranking barcodes by their total UMI counts and visually selecting a knee point, where a steep dropoff in counts occurs^3,12^. Droplets with higher counts are expected to contain cells or nuclei, whereas droplets with lower counts are expected to contain ambient RNA. However, a clear separation between the two may not occur, especially if the amount of debris is high and the droplet RNA content is low, as we show is the case with frozen solid tissues. Additionally, an *ad hoc* cutoff of the percent of reads originating from the mitochondria (a measure of extranuclear contamination) can help to filter droplets^12^. Again, the choice of a cutoff may be arbitrary or unclear. The recent method EmptyDrops addresses this filtering issue for scRNA-seq by estimating a Dirichlet-Multinomial distribution of the ambient RNA. It then removes droplets by testing if their expression profile deviates significantly from the ambient profile using a Monte Carlo approach^12^. However, while this works for single-cell, we show that these methods underperform when applied to snRNA-seq.

Here we show that, in snRNA-seq, using a hard cutoff to remove droplets can result in a substantial loss of nuclear droplets and inclusion of debris droplets. Importantly, we demonstrate that including these contaminated droplets can lead to spurious clustering and false positive cell types. To overcome this, we built a fast filtering pipeline that uses a likelihood-based approach to model debris and cell type RNA distributions with a multinomial distribution. The parameters of the model are inferred using semi-supervised EM^13,14^, where all droplets below a hard count threshold are fixed as debris. Then, droplets above the threshold are classified as debris or nucleus based on its likelihood in the inferred model. This approach has been successfully applied to the information retrieval and text mining fields^15^. Similar to reads, word occurrences in a document can be modeled with a multinomial distribution, and documents can belong to separate topics, leading to a mixture model.

We developed this pipeline into an approach, termed Debris Identification using EM (DIEM), which robustly removes background droplets from both scRNA-seq and snRNA-seq data. In contrast to hard count and EmptyDrops filtering, DIEM takes into account multiple cell types when modeling gene expression distributions. This resulted in more accurate filtering and higher quality clustering of snRNA-seq data, particularly when applied to frozen tissue. Filtering of scRNA-seq data using DIEM also led to inclusion of more cell-containing droplets and higher resolution in clustering.

## Results

### snRNA-seq produces clusters driven by high amounts of background RNA contamination

Isolation of nuclei for snRNA-seq relies on lysis of the cell membrane, releasing cytoplasmic RNA, in addition to cell-free RNA, into the solution. This extranuclear RNA can become encapsulated into droplets, with or without nuclei, and lead to biases in downstream analysis; particularly, it may lead to spurious or contaminated cell-types in downstream clustering. We evaluated the extent of contamination and its effect on clustering in three distinct snRNA-seq data sets: 1. *in vitro* differentiating human preadipocytes (DiffPA) (n=1), 2. freshly dissected mouse brain tissue (n=1), and 3. frozen human subcutaneous adipose tissue (AT) (n=6).

We initially ran a clustering analysis in the three data sets by filtering out droplets with a hard-count threshold^3,8–11^. This threshold can be selected manually, as the knee point^3^, or by dividing the total count of the 99% quantile of expected cells by 10^16^. Since we observed that the knee point could not be reliably estimated or was not evident in the AT samples (Fig. 1a), we used the quantile-based threshold for further analyses. We evaluated the extent of extranuclear contamination on clustering by quantifying the percentage of mitochondria-derived UMIs (MT%). We chose to use mitochondrial RNA as a measure extranuclear RNA contamination because it is one of the only true sources of background RNA and is present in all snRNA-seq data sets. However, we note that other sources of extranuclear RNA can exist. Hemoglobin mRNA, which is predominantly expressed in erythrocytes, can also serve as another negative control for tissues where blood is present^17^.

**Figure 1.**
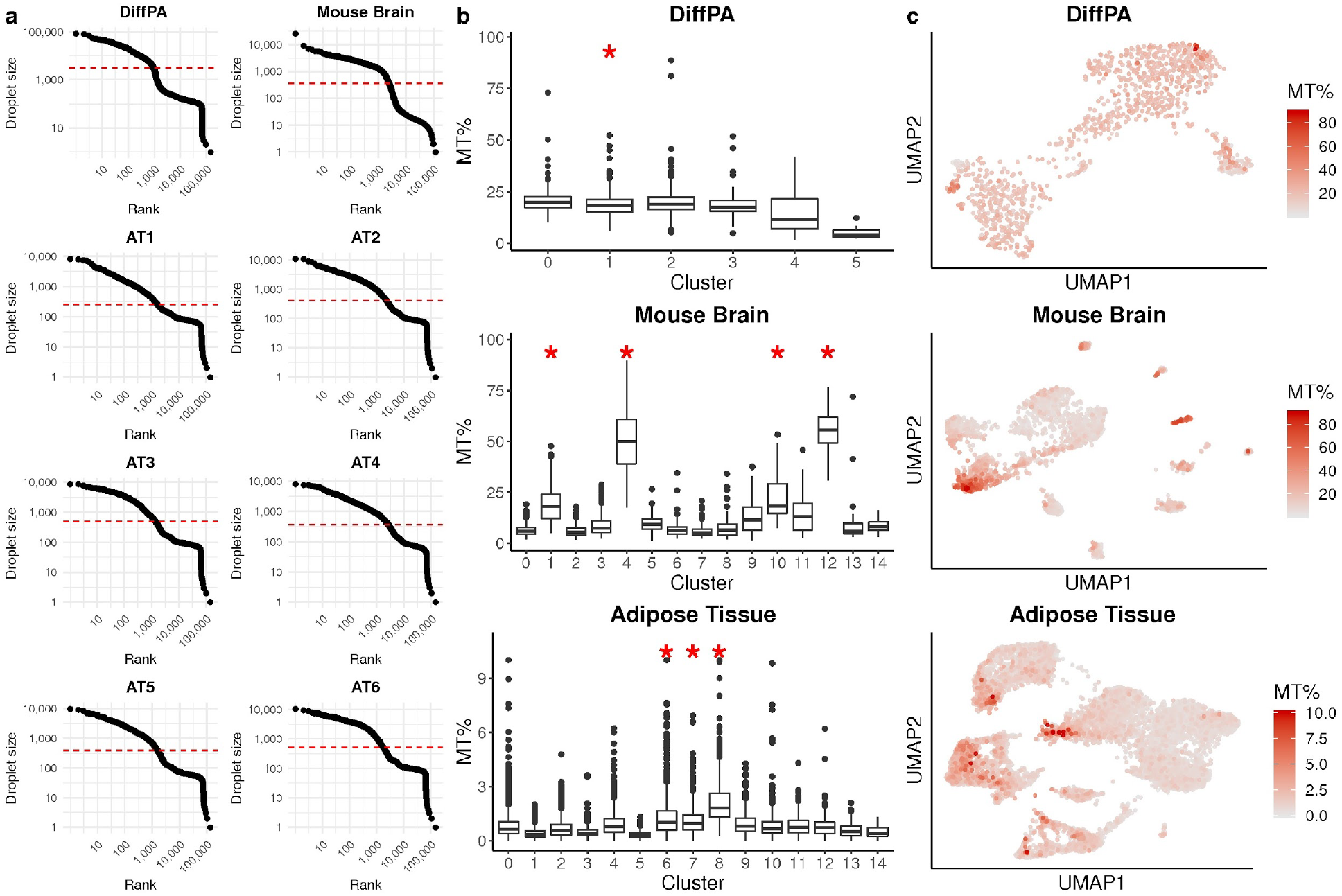
Applying a hard count threshold fails to remove droplets contaminated with background RNA in snRNA-seq. **a**, Barcode-rank plots showing the droplet size (the total number of UMI read counts) of each droplet in descending order for the differentiating preadipocytes (DiffPA), mouse brain, and six human frozen adipose tissue (AT) snRNA-seq samples. The dotted red line indicates the quantile-based threshold. **b**, The percent of UMIs mapping to the mitochondrial genome (MT%) for each droplet are shown in a box plot, separated by clusters identified by Seurat^19^. The red asterisk shows which clusters contain a significant (FDR < 0.05) MT-encoded gene as an up-regulated marker when compared to droplets in all other clusters. **c**, UMAP^22^ visualization after quantile-based filtering shows clustering of droplets with a high MT%.

To test whether the quantile-based method could effectively remove debris-contaminated nuclei, we investigated the relationship between MT% and the total number of UMIs in a droplet. As expected, we observed that nuclei below the threshold tended to have more droplets with higher levels of mitochondrial contamination (Supplementary Fig. 1). However, we noticed that some droplets above the threshold also contained high MT% similar to that of background RNA-enriched droplets below it. For example, in the mouse brain data set, the mean MT% of droplets below and above the quantile-based threshold was 56.6% and 23.0%, respectively. Still, 16.3% of droplets above the quantile-based threshold had an MT% greater than the mean of droplets below it. Additionally, each data set also contained a smooth range of MT% across droplets, suggesting difficulties in selecting a hard cutoff to remove contamination (Supplementary Fig. 1). Using mitochondrial contamination as a proxy, our results imply that extranuclear RNA contamination can affect droplets in higher UMI ranges above typical hard count thresholds.

Clustering of droplets based on gene expression profiles is used to identify cell types present in a heterogeneous source of cells such as tissues^18^. Since background RNA contamination affects the gene expression profile of a droplet, we investigated whether this could cause spurious clustering and false-positive cell types driven by this background RNA. For the DiffPA, mouse brain, and AT data sets, we took the filtered barcodes from the commonly applied quantile-based threshold and clustered them using Seurat^19^. We checked whether mitochondrial genes were up-regulated in any of these clusters, which is indicative of background RNA contamination. In all three data sets, we observed at least one cluster containing significantly higher UMI counts from MT-encoded genes (Fig. 1b). Dimensionality reduction and visualization with UMAP^20^ showed that droplets with higher levels of mitochondrial RNA tended to cluster together (Fig. 1c). This problem was more apparent for the tissue data sets (mouse brain and human adipose) than for the *in vitro* (DiffPA) experiment. Overall, in the DiffPA, mouse brain, and human AT snRNA-seq data sets, using a hard count threshold failed to remove droplets contaminated with extranuclear RNA and resulted in clusters marked by high MT%.

### Nuclear and debris droplets demonstrate distinct RNA profiles

Since a droplet’s total UMI counts did not effectively discriminate nuclei from debris, we postulated that the expression profile of a droplet could be used to differentiate them if there were sufficient differences in RNA abundance between cell types and debris. Specifically, we hypothesized that there would be nuclear-encoded genes that show differential abundance. Thus, we evaluated the extent of differences between the debris and nuclear RNA profiles. We separated droplets into debris- and nuclear-enriched groups using a threshold of 100 total UMI counts. Although a large amount of droplets above 100 UMI counts consist of debris and would lead to a loss of power, we use this threshold to ensure that no droplets below it contain nuclei. We evaluated the difference between the debris and nuclear RNA profiles by running a paired differential expression (DE) analysis in the six human AT samples. Of 19,934 genes detected, 3,417 (17.1%) were DE between the nuclear- and debris-enriched groups at a Bonferroni-adjusted p-value threshold of 0.05 (Fig. 2a). To see if these differences were preserved across the DiffPA, mouse brain, and six AT data sets, we correlated the nuclear vs. debris log fold changes of the genes in common. Among the 8,924 genes expressed in all three data sets, we found that all log fold changes were significantly correlated (p<2.2×10^−16^) across all pairs (mean *R* = 0.64), with the human data sets showing the highest correlations (Supplementary Fig. 2).

Since the nuclear-enriched group is not homogeneous, but rather originates from distinct cell types with different RNA distributions, we also looked at differences between the debris group and cell types. In addition, we compared the cell type-debris differences with the cell type-cell type differences. Using the six AT samples, we ran a paired DE analysis between the cell types and debris droplets (total UMI counts < 100). Among 14 debris-cell type pairs, the average percent of genes that are DE was 5.8% (Fig. 2b). We then compared this to the DE between a cell type and all other cell types. Among these 14 pairs, the average percent of genes DE between cell types was slightly lower at 4.5% (t-test p=0.23; Fig. 2b,c). Overall, we found significant differences between debris and nuclei RNA profiles, and that the differences between debris and cell types were within the same order of magnitude as the cell type-cell type differences.

**Figure 2.**
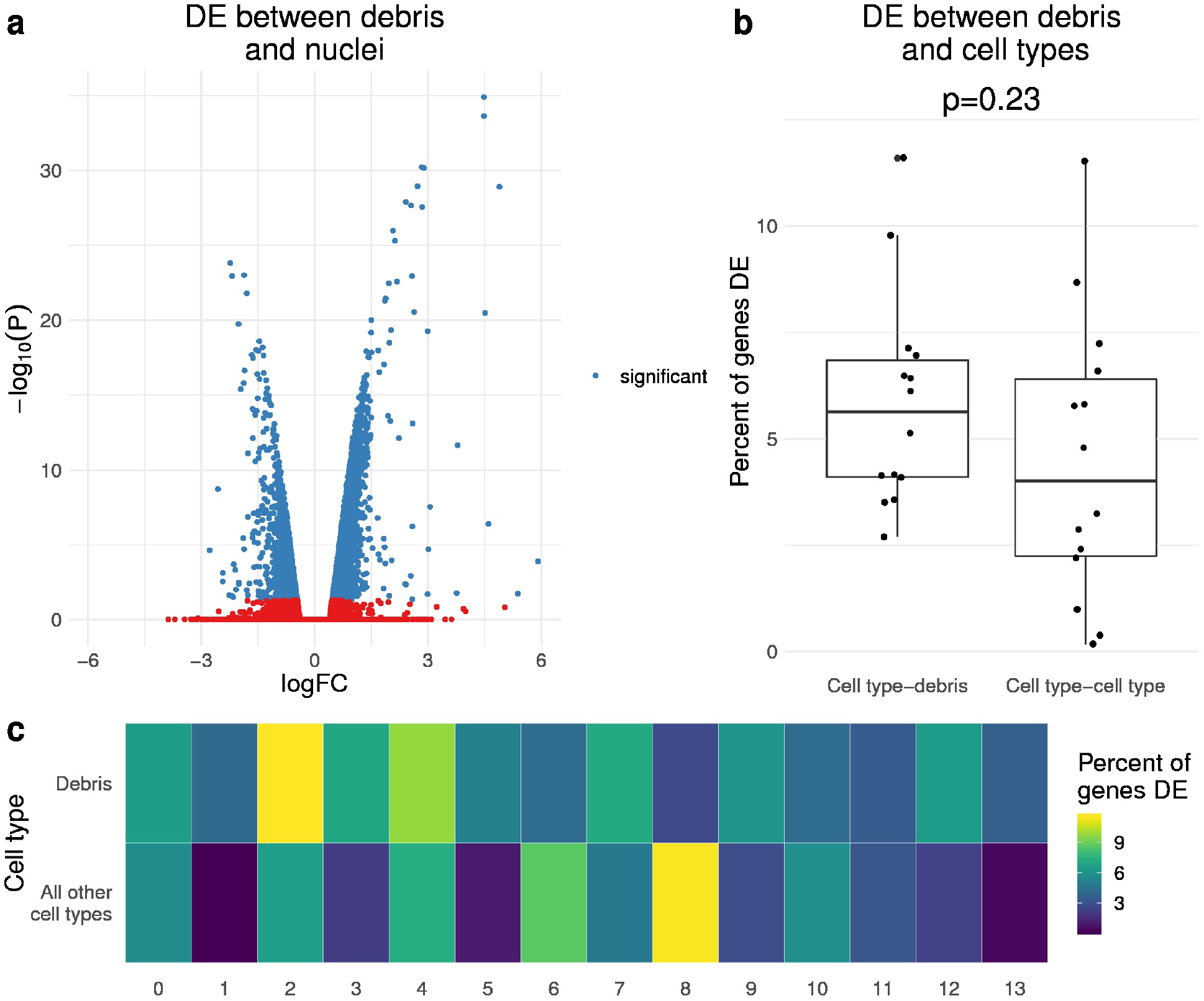
Debris-containing and nuclei-containing droplets show distinct gene expression profiles. **a**, Differential expression (DE) between droplets with less than 100 UMI counts (debris) and greater than or equal to 100 UMI counts (nuclei) in the 6 human adipose tissue (AT) samples. The volcano plot shows the log fold change on the x-axis and negative log transformed p-value on the y-axis. The genes colored in blue are DE with a Bonferroni-corrected p-value < 0.05. A positive log fold change indicates over-expression in the debris group. **b**, DE between debris droplets and cell types in the 6 human AT samples. Cell types are estimated from clustering droplets filtered after quantile-based thresholding. A box plot shows the percent of expressed genes that are DE (Bonferroni p<0.05) between a cell type-debris pair, and a cell type-cell type pair. The p-value was calculated from a student’s t-test between cell type-debris percents and cell type-cell type percent. **c**, The heatmap shows the percent of expressed genes that are DE between the given pairs.

### Overview of a novel EM-based approach to cluster and remove debris droplets from snRNA-seq data

Since we observed differences in RNA abundance between cell types and debris, we developed an approach to remove debris-containing droplets based on the distribution of read counts. Our approach uses a multinomial mixture model for *k* + 1 mixtures, corresponding to *k* cell types and 1 debris group. To estimate the parameters of the multinomial mixture model, we run semi-supervised expectation maximization^13,14^ by fixing droplets that fall below a hard count threshold as debris. After fitting the model, we assign droplets above the threshold to their group of origin (debris or a cell type) based on their posterior probability. Figure 3a shows an overview of this model. We termed this method Debris Identification using Expectation Maximization (DIEM). We compared our approach with the quantile-based method and the EmptyDrops method in the DropletUtils package^12^.

**Figure 3.**
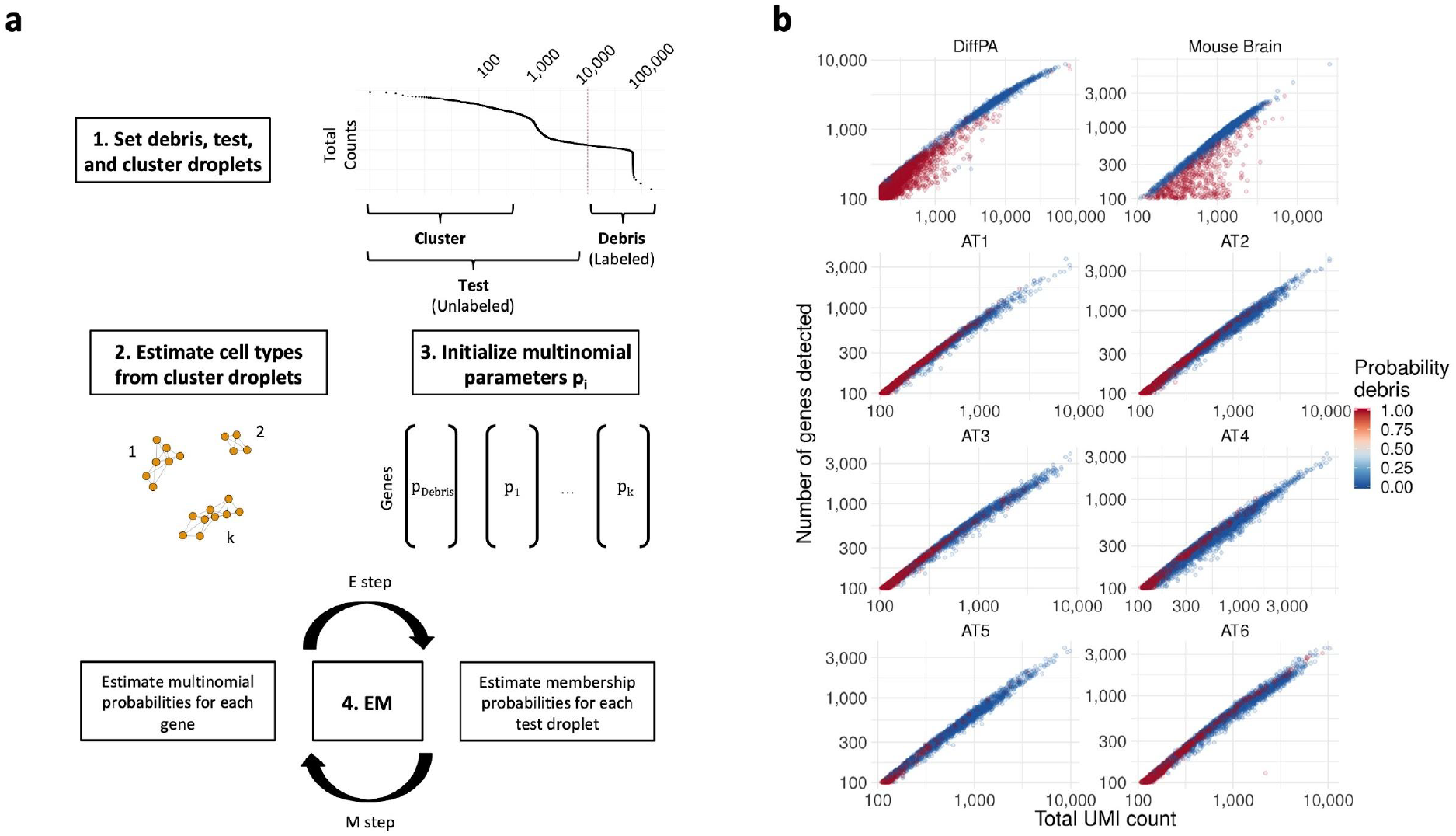
EM-based approach estimates parameters of debris and nuclear distributions and removes contaminated droplets. **a**, Overview of DIEM approach to remove debris-contaminated droplets. Expectation Maximization (EM) is used to estimate the parameters of a multinomial mixture model consisting of debris and cell type groups. The label assignments of droplets below a pre-specified threshold are fixed to the debris group, while the test set droplets above this rank are allowed to change group membership. The cell types are estimated by clustering the top *N* droplets. The means *p*_*i*_ of the debris and cell type multinomials are initialized from the droplets in the debris and initialized cell type groups, respectively. In the E step, the posterior probabilities of the groups are estimated for the unlabeled droplets, keeping the labeled droplets fixed to the debris group. In the M step, the model parameters are re-estimated. The algorithm converges until the percent change in likelihood is smaller than a pre-specified value. **b**, Scatterplots of droplets from snRNA-seq of the differentiating preadipocytes (DiffPA), mouse brain, and human frozen adipose tissue (AT) data sets, with total UMI counts on the x-axis and total number of genes detected on the y-axis. Droplets are colored by the DIEM-assigned posterior probability that it originated from a debris-contaminated cluster. Those in red are called as debris while the blue droplets are called as nuclei.

We first looked at the characteristics of the droplets removed by DIEM in the adipose tissue data set. We observed that DIEM removed droplets across a range of total UMI counts (Fig. 3b). Of a total of 15,855 test droplets, DIEM removed 1,893 (11.9%). In comparison, the quantile-based approach removed 4,524 (28.5%) while EmptyDrops removed 4,353 (27.5%). Next, we asked whether these removed droplets were truly contaminated or whether they solely contained nuclei. We clustered the removed droplets and compared their characteristics to clusters identified from the passing droplets. UMAP visualization showed that the droplets removed by the EmptyDrops and quantile-based methods formed more distinct clusters, suggesting that these droplets originated from cell types (Supplementary Fig. 3). In addition, we looked at the distribution of MT% in these removed clusters. Some of the clusters formed by the EmptyDrops- and quantile-removed droplets showed low levels of MT% similar to that of the clusters that passed filtering, suggesting they are not contaminated with extranuclear RNA (Supplementary Fig. 3). In contrast, the DIEM-removed clusters had a higher MT% than the majority of the clusters that passed filtering. The average MT% of the DIEM-removed droplets was 2.2%, while the averages of the EmptyDrops and quantile thresholding ones were 1.2% and 1.4%, respectively. Overall, these results suggest that DIEM preferentially removes MT-contaminated droplets.

The incorporation of cell types should result in a more realistic model of the snRNA-seq data. DIEM implicitly assigns each droplet to a cell type after fitting the mixture model with EM. To evaluate the accuracy of these assignments, we investigated the DIEM-assigned clusters using a principal component analysis (PCA). We observed that the DIEM-assigned clusters tended to aggregate in the PCA plots, particularly in the PC 1-2 subspace (Supplementary Fig. 4). Additionally, we evaluated how well the multinomial clusters corresponded to clusters that would be identified from Seurat. While the number of cell types estimated by DIEM was fewer than the number estimated by Seurat, the Seurat clusters overlapped highly with the DIEM clusters (mean percent overlap 88.8%; Supplementary Fig. 4). Together, these results suggest that DIEM leverages cell type heterogeneity by accurately modeling the transcriptomes of major cell types.

### DIEM produces cleaner clusters containing less extranuclear RNA contamination

After verifying that DIEM preferentially removes debris-contaminated droplets, we evaluated its ability to produce valid clusters. We ran a Seurat^19^ pipeline to obtain clusters following each of the three filtering methods. Then, we overlapped the resulting clusters to identify cell types that were removed and/or added by DIEM. In all three data sets, DIEM preserved most cell types identified from both the quantile and EmptyDrops approaches while removing clusters that were characterized by high levels of extranuclear RNA contamination (Fig. 4a,b). We also noticed that DIEM removed a low MT% cluster from the DiffPA dataset. This cluster, identified by both the quantile and EmptyDrops methods, contained much lower levels of the nuclear-localized lincRNA MALAT1^21^ and lower total UMIs, making it unclear whether these droplets contain nuclei (Supplementary Fig. 5). In some instances, we noticed that DIEM filtering resulted in the merging of two cell types. We asked whether the cell type was removed by our method or was merged because of changes to the input of the clustering algorithm. We increased the clustering resolution parameter of Seurat^19^ to more sensitively detect a larger number of more related clusters. Increasing this parameter in the DIEM-filtered data sets split the previously merged clusters and showed a one-to-one correspondence with the quantile-based and EmptyDrops clusters (Supplementary Fig. 6). Overall, we found that DIEM filtering preserves cell types identified by previously established methods.

**Figure 4.**
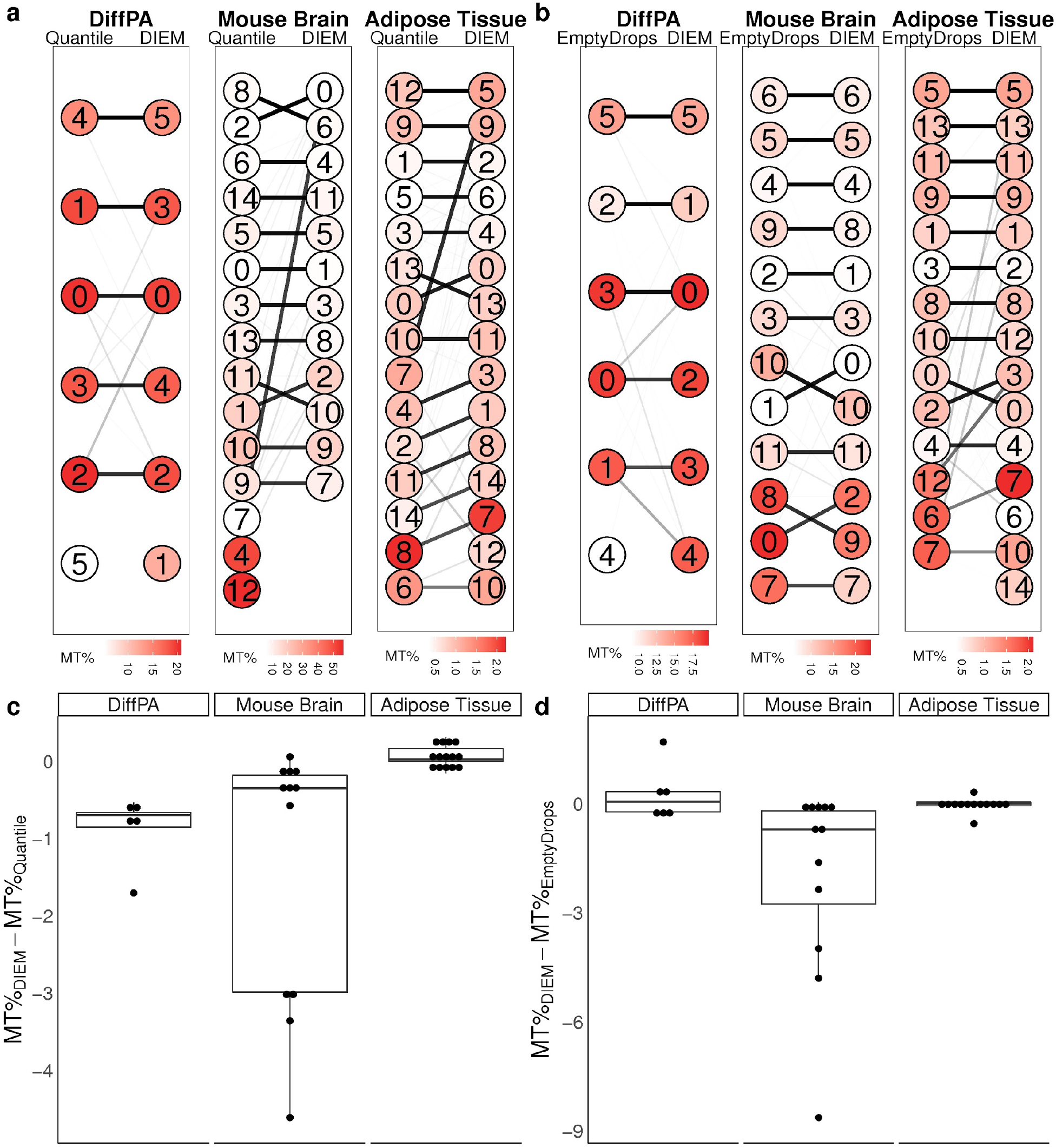
DIEM removes contaminated clusters while preserving cell-types identified by previous methods. **a,b**, Graphs showing the overlap of clusters for the differentiating preadipocytes (DiffPA), mouse brain, and human frozen adipose tissue (AT) data sets. Each circle is a cluster identified after filtering with the method specified at the top. The thickness of the line between a cluster pair indicates the percent of droplets from the **(a)** quantile or **(b)** EmptyDrops^12^ cluster that are shared with the connected DIEM cluster. No lines emerging from an EmptyDrops cluster means that droplets from that cluster are removed by DIEM. The color of each cluster indicates the average percent of UMIs mapping to the mitochondrial genome (MT%). **c,d**, Bar plots showing the difference in MT% after filtering using DIEM when compared to the **(c)** quantile or **(d)** EmptyDrops methods. Each dot represents a cluster pair corresponding to the two filtering methods. A negative difference indicates a decrease in MT% after filtering with DIEM in that cluster.

Next, we quantified the extent to which DIEM was able to remove extranuclear RNA contamination within individual clusters. By comparing corresponding clusters identified by the compared methods, we found that, on average, DIEM decreased the average MT% of cluster-specific droplets, especially in the mouse brain data set (Fig. 4c,d). After observing a MT% decrease within clusters, we also compared the total number of statistically significant MT marker genes. In the DiffPA data set, the quantile method yielded 3 MT cluster markers (Supplementary Fig. 7). In both the mouse brain and AT data sets, DIEM filtering resulted in the fewest number of total MT markers (Supplementary Fig. 7). This suggests that DIEM filtering results in less clustering of extranuclear RNA contamination. UMAP visualization^22^ also reflected this reduction in MT%-driven clustering after DIEM filtering (Fig. 5). To conclude, our filtering approach produced clusters containing as much or less extranuclear RNA contamination (assessed by MT%) than those produced by the quantile or EmptyDrops methods.

**Figure 5.**
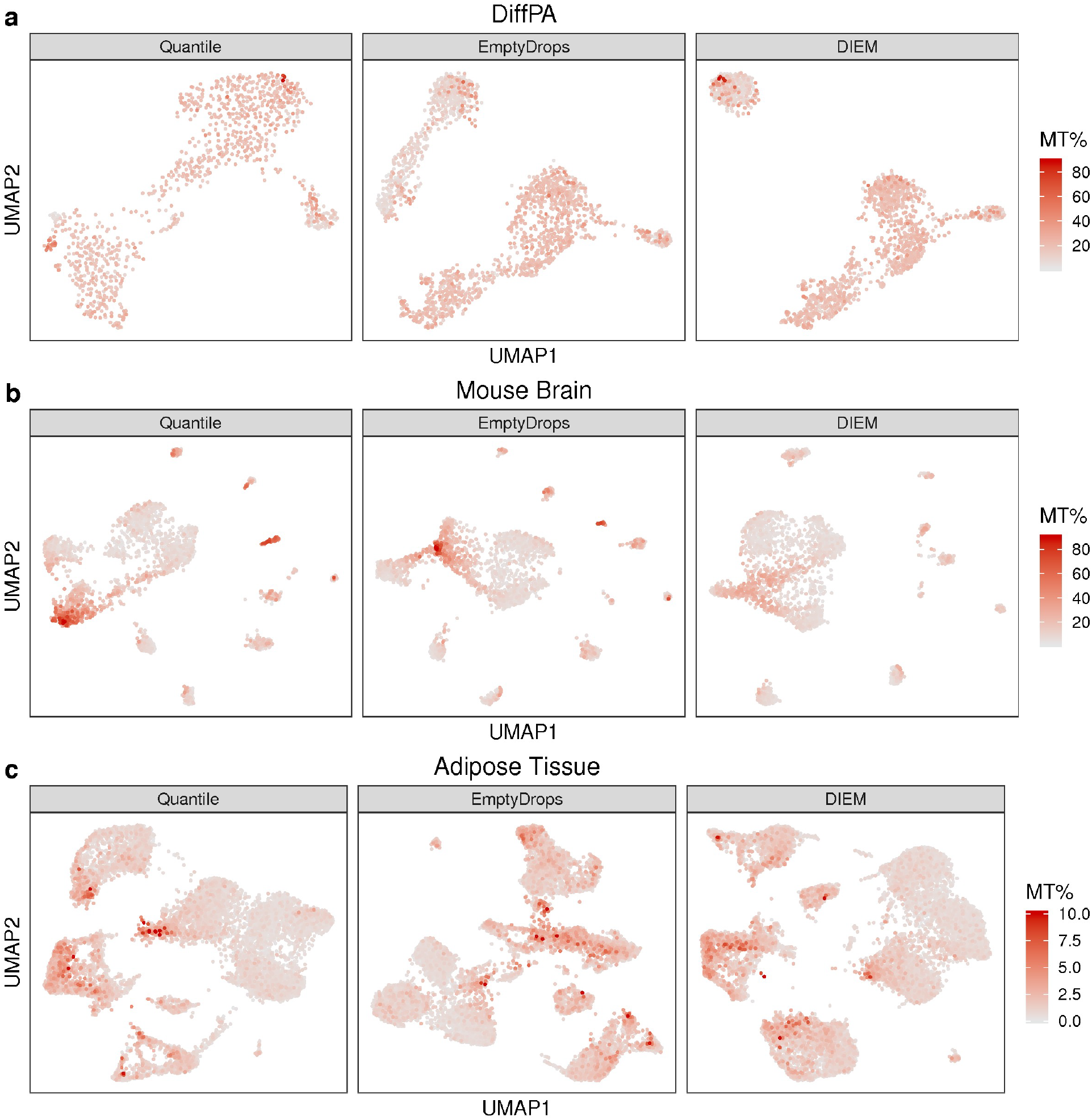
DIEM filtering results in less mitochondrial RNA contamination. **a,b,c**, UMAP^22^ visualization of droplets for each filtering method in the **(a)** differentiating preadipocytes (DiffPA), **(b)** mouse brain, and **(c)** human frozen adipose tissue (AT) data sets. Droplets are colored by percent of UMIs mapping to the mitochondrial genome (MT%).

### DIEM filtering removes debris from single-cell RNA-seq

In addition to filtering snRNA-seq, we also investigated whether our approach could be applied to single-cell RNA-seq data. To this end, we ran DIEM on ~68,000 PBMCs from blood^16^. Similar to the snRNA-seq datasets, we compared DIEM to the quantile and EmptyDrops^12^ methods. DIEM and EmptyDrops kept 69,521 and 69,981 droplets, respectively, while the quantile threshold kept 66,233. We then investigated the characteristics of the 460 droplets removed by DIEM and kept by EmptyDrops. We found that these 460 droplets tended to have higher MT%, as well as a higher percentage of UMIs aligning to MALAT1 (Fig. 6a). Although these metrics no longer serve as negative and positive controls as they do in snRNA-seq, they are consistent with a ruptured cell membrane. This suggests that EmptyDrops retains droplets with dying cells whereas DIEM removes them. We next compared the resulting clusters from each of the three methods. While the quantile approach identified 17 clusters, both DIEM and EmptyDrops identified 19 (Fig. 6b-d). The droplets in these two additional clusters were present in the quantile data set, however, it is likely that the increased number of cells resulted in better resolution to identify more cell types with DIEM and EmptyDrops. Furthermore, there was a high concordance between the cell types identified with EmptyDrops and those identified with DIEM (Fig. 6e,f). These results indicate that our filtering approach can be applied to identify cell-containing droplets in single-cell RNA-seq data.

**Figure 6.**
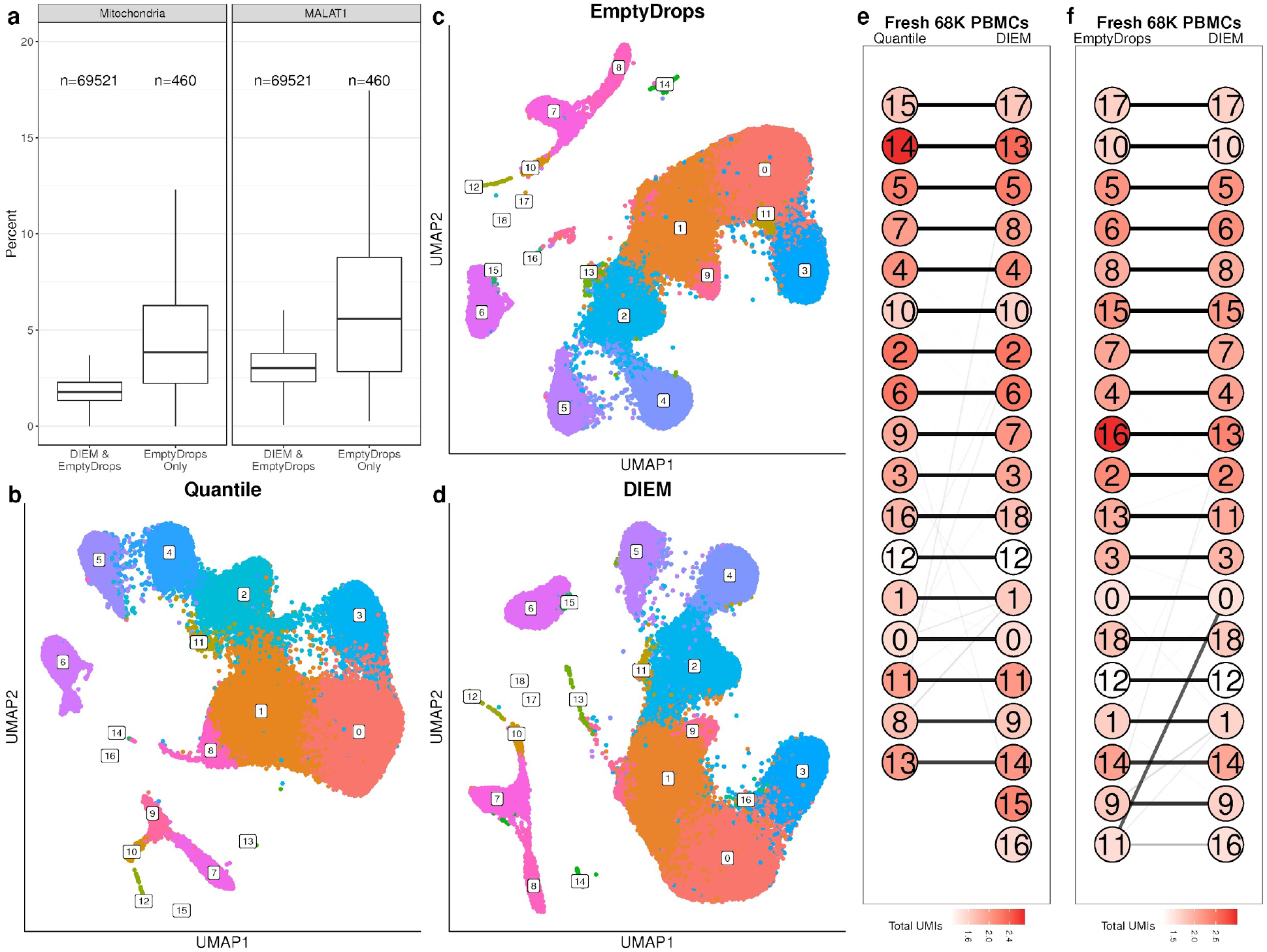
DIEM filtering identifies additional droplets in single-cell RNA-seq of fresh PBMCs. **a**, Boxplots showing the percent of unique molecular indices (UMIs) mapping to the mitochondria (left) and the percent of MALAT1 UMIs (right) in the fresh 68K PBMC data set^16^. The DIEM and EmptyDrops^12^ set includes the droplets identified by both DIEM and EmptyDrops, while the EmptyDrops Only set includes those droplets removed by DIEM and kept by EmptyDrops. **b,c,d**, UMAP^22^ visualizations of clustering after **(b)** quantile, **(c)** EmptyDrops, and **(d)** DIEM filtering. **e,f**, Concordance of clusters identified between **(e)** quantile and DIEM filtering, and **(f)** EmptyDrops and DIEM filtering. The thickness of a connecting line between a quantile/EmptyDrops cluster and a DIEM cluster represents the percent of droplets in the quantile/EmptyDrops cluster that are contained in the connecting DIEM cluster. No lines emerging from a quantile/EmptyDrops cluster means that droplets from that cluster are removed by DIEM. The color of a cluster circle indicates the average percent of mitochondrial UMIs per cluster (MT%).

## Discussion

The snRNA-seq approach is an adaptation of scRNA-seq that allows for cell-type identification when isolation of a cell suspension is not possible, such as in frozen tissues. We have shown here that snRNA-seq is subject to background contamination from extranuclear RNA and that it can drive spurious clusters and false positive cell-types if not properly accounted for. We also show that current methods, such as the commonly applied hard count threshold, do not effectively address this problem in snRNA-seq. To this end, we searched for and found differences in the gene expression profiles from the debris and cell types. This motivated us to use the RNA profile of a droplet for filtering. We developed an approach, termed DIEM, which uses a multinomial mixture model and then estimates the probability that a droplet originates from either the debris or a cell type. We found that DIEM removes debris-contaminated droplets from both snRNA-seq and scRNA-seq data, leading to removal of spurious cell-types while preserving biologically plausible cell-types.

Even though snRNA-seq recovers less RNA than scRNA-seq and thus retrieves less information about cell types, there are advantages to using nuclei over cells. snRNA-seq has been shown to reduce dissociation biases present in scRNA-seq, leading to more accurate profiling of cell types in tissue^23^. Another important reason to use snRNA-seq is that scRNA-seq may be practically impossible. This can occur with frozen tissues, since thawing cells can lyse the outer membranes and preclude a suspension of single cells required for droplet-based technologies^3^. This prevents the application of scRNA-seq to biobanked snap-frozen human tissues. In order to leverage existing, phenotyped human datasets with biobanked tissues, snRNA-seq may be the only viable option to profile cell types. As we have shown, however, snRNA-seq of frozen tissue results in contamination of droplets across a large range of UMI counts, making it difficult to use a hard count threshold to remove background debris. Even from fresh tissue and cells, we still observed downstream clusters affected by the extranuclear RNA. Therefore, we expect DIEM to help produce cleaner snRNA-seq data sets from a variety of input sources, but especially from frozen tissues.

When compared to the EmptyDrops method^12^, we found that DIEM can better remove extranuclear RNA contamination and retain a higher number of nuclei in snRNA-seq data. EmptyDrops, however, was originally developed and tested on single cell data and thus, the assumptions behind the model are different than that of DIEM. EmptyDrops assumes a single Dirichlet-multinomial distribution for the background RNA, and uses Monte Carlo sampling to determine how significant the deviation of a droplet is from it. It also safeguards from removing cell-types that are similar to the background by assuming that all droplets above a calculated knee point are true cell-containing droplets. Our approach addresses this issue directly by modeling the cell types present in the mixture. We have shown that the difference between the cell types and the debris are within the same order of magnitude as the differences between the cell types, highlighting the need to account for heterogeneity. Accordingly, DIEM removed fewer droplets from putative cell types when assessed by clustering filtered out droplets.

We focused the application of our approach on snRNA-seq data in this paper because there is a pressing need for filtering in these data sets with lower RNA content. In single-cell RNA-seq, the higher RNA content of cells typically allows the total UMI count of a droplet to serve as a sufficient discriminator between debris and cells^3^, although this may not always be the case^12^. However, our approach simply requires that there are differences in the distributions between the debris and the cell types. As this can also occur in single cell RNA-seq data, we applied DIEM to the 10X 68k PBMC data set^16^ (n=1 individual) and found that the resulting clusters were concordant with previous filtering methods. Thus, DIEM can also be used to filter single cell RNA-seq data. We expect our expression-based filtering to be more useful for tissues where cell-free RNA^24,25^, hemoglobin from lysed red blood cells, or total RNA from a diversity of lysed cells are more prominent.

Running scRNA-seq on fresh human tissue at a large scale may be prohibitively difficult considering the requirement to immediately process a fresh biopsy for scRNA-seq. Therefore, snRNA-seq of frozen tissues offers a viable alternative to process samples at a higher throughput. Our method was designed to computationally remove background debris contamination from snRNA-seq data of frozen tissues. We expect that DIEM will enable the analysis of a larger number of samples from frozen tissue snRNA-seq data, thereby removing the need to coordinate the acquisition of fresh tissue samples and processing of single cell libraries.

## Methods

### Single-nucleus RNA-seq of human subcutaneous adipose tissue, differentiating preadipocytes, and mouse brain

Frozen subcutaneous adipose tissue was processed separately for each of the 6 samples. Each of the 6 participants gave a written informed consent. The study protocol was approved by the Ethics Committee at the Helsinki University Hospital, Helsinki, Finland. Tissue was minced over dry ice and transferred into ice-cold lysis buffer consisting of 0.1% IGEPAL, 10mM Tris-Hcl, 10 mM NaCl, and 3 mM MgCl2. After a 10 minute incubation period, the lysate was gently homogenized using a dounce homogenizer and filtered through a 70 μm MACS smart strainer (Miltenyi Biotec #130-098-462) to remove debris. Nuclei were centrifuged at 500 g for 5 minutes at 4°C and washed in 1 ml of resuspension buffer (RSB) consisting of 1X PBS, 1.0% BSA, and 0.2 U/μl RNase inhibitor. We further filtered nuclei using a 40 μm Flowmi cell strainer (Sigma Aldrich # BAH136800040) and centrifuged at 500 g for 5 minutes at 4°C. Pelleted nuclei were re-suspended in wash buffer and immediately processed with the 10X Chromium platform following the Single Cell 3’ v2 protocol. After library generation with the 10X platform, libraries were sequenced on an Illumina NovaSeq S2 at a sequencing depth of 50,000 reads per cell. Reads were aligned to the GRCh38 human genome reference with Gencode v26 gene annotations^26^ using the 10X CellRanger 2.1.1 pipeline. A custom pre-mRNA reference was generated to account for unspliced mRNA by merging all introns and exons of a gene into a single meta-exon.

We obtained and cultured the primary human white preadipocyte cells as recommended by PromoCell (PromoCell C-12731, lot 395Z024) for preadipocyte growth and differentiation into adipocytes. Cell media (PromoCell) was supplemented with 1% penicillin-streptomycin. We maintained the cells at 37ºC in a humidified atmosphere at 5% CO2. On day 6 of differentiation, we rinsed the cells with 1x PBS and added ice-cold lysis buffer (3 mM MgCl2, 10 mM Tris-HCl, 0.5% Igepal CA-630, 10 mM NaCl). The cells were gently scraped from the plate and centrifuged at 500 x g for 5 minutes at 4ºC. Nuclei were washed with 1 ml of resuspension buffer (RSB; 1% BSA, 100 μl RNase inhibitor in 1x PBS) and centrifuged again to remove cellular debris. After the second centrifugation, nuclei were washed with 1 ml RSB and filtered through a 40 μm filter. Cells were counted, then centrifuged again and resuspended in the proper volume of RSB to obtain 2000 nuclei/μl. The 10X library preparation, sequencing, and data processing were done using the same protocol as for the adipose tissue.

For the mouse brain data, we downloaded the raw UMI count data matrix from the 10X website. The data set titled “2K Brain Nuclei from an Adult Mouse (>8 weeks)” was downloaded from https://support.10xgenomics.com/single-cell-gene-expression/datasets/2.1.0/nuclei_2k. The 10X human 68K PBMC data were downloaded from

### Filtering droplets using a quantile threshold and EmptyDrops

Common methods for removing debris from snRNA-seq data rely on using a hard count threshold^3,8–11^. In the three data sets, we applied a quantile-based cutoff, similar to that implemented by the 10X CellRanger software. Droplets are ranked in decreasing order of total counts. The 99th percent quantile of the top *C* barcodes of total counts is divided by 10 to obtain the threshold *T*, where *C* is 3,000 for our analyses^16^. The 99th percentile is used to exclude any doublets from the derivation. Droplets with greater than or equal to *T* counts were included as nuclei. For comparison with EmptyDrops^12^, we ran the method using default parameters. EmptyDrops calculates a Monte Carlo p-value that gives the probability that a droplet’s expression profile is the same as that of the ambient RNA. We removed droplets with a false discovery rate (FDR) q value greater than 0.05.

### Differential expression between nuclear-enriched and debris-enriched droplets

To identify genes differentially expressed (DE) between the background-enriched and nuclear-enriched groups, we set a hard count threshold to naively assign droplets to either group. Droplets with total UMI counts below 100 and greater than or equal to 100 were assigned to the background-enriched and nuclear-enriched groups, respectively. This ensures that the majority of droplets containing nuclei are found in the nuclear-enriched group. For each gene, reads were summed across all droplets in each of the two groups to estimate the RNA profiles. Read counts were normalized using trimmed mean of M-values (TMM) as implemented in edgeR^27,28^. For identifying differentially expressed genes, we used a paired design with the six adipose tissue samples by treating the background-enriched and nuclear-enriched counts of an individual as a paired sample (total n=12). We then used the edgeR package^27,28^ to run differential expression. We only kept genes with a counts per million (CPM) of greater than 0 in at least 6 of the 12 groups. Next, we used the estimateDisp function to estimate the dispersion with the paired design matrix. The quasi-likelihood fit and F test functions glmQLFit and glmQLFTest were used to calculate statistical significance. We adjusted for multiple testing using a Bonferroni-corrected p-value threshold of 0.05.

To identify DE genes between the debris and cell types, we used the clusters identified after quantile-based filtering to approximate the cell types. For each of the six samples, we subsampled the debris droplets (with total UMI counts less than 100) to 9,000 droplets to obtain a similar read depth as contained in the cell type groups. For the debris and cell type groups, reads were summed across the corresponding droplets to obtain the RNA profile used as input. Differential expression was performed by comparing debris vs. cell type or cell type vs. all other cell types using a paired design. The filtering and analysis was performed in the same manner as the debris vs. nuclear DE analysis above.

### DIEM Algorithm

Our filtering approach models droplet-based single-cell or single-nucleus data with a multinomial mixture distribution. Read counts in a droplet follow a multinomial with parameters conditional on whether it contains a cell type or debris. However, the parameters of the multinomials and droplet assignments are unknown. Therefore, we estimate the parameters of the model using semi-supervised expectation maximization (EM)^13,14^. Following this, droplets are classified as nuclei or debris according to their likelihood. To obtain the number of cell types *k* and to reasonably initialize them for EM, we cluster the high-count droplets that are expected to contain mostly nuclei or cells.

In more detail, let *X* denote a *g* x *n* matrix containing the read/UMI counts from a single-cell or single-nucleus data set with *g* genes and *n* droplets. We include droplets with at least 1 read/UMI count. Our goal is to assign the *n* droplets into one of *K* groups (*K* - 1 cell types and debris). We define *x*_*i*_ as the *i*th column of *X* giving the counts of droplet *i*. We model the read counts *x*_*i*_ with a multinomial distribution, with the gene probabilities *m*_*k*_ = *p_1,k_*, …, *p_G,k_* conditional on group *k* ∈ {1,..., *K*}. In addition to the observed read counts *x*_*i*_, droplet *i* has the unobserved latent variable *z_i_* ∈ {1,..., *K*} that describes the cell type or debris origin of the droplet. The log-likelihood of the data is therefore:

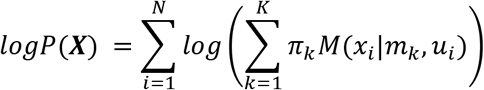

Here, *u*_*i*_ is the total number of read/UMI counts in droplet *i*, *m*_*k*_ contains the gene probabilities for group *k*, *π_k_* is the mixture coefficient for group *k*, and *M* denotes the probability mass function of the multinomial distribution. Since an analytical solution cannot be derived based on the likelihood of the model, we use a semi-supervised EM algorithm to estimate the parameters *m_1_*, …, *m*_*k*_ and *π_1_*, …, *π_k_* by maximizing the expected complete data log-likelihood using the latent variables {*z_i_*}. Before running EM, we estimate the number of cell types and initialize the parameters of multinomial mixture model as described below.

First, we assign droplets into the test set, the debris set, and the cluster set. The test set consists of the droplets we would like to classify, while the debris set consists of droplets that are very likely to contain debris. We define the test set as those droplets above a count threshold *T*, where *T* is the maximum of 100 or the total counts of the *U*th size-ranked droplet. In this paper, we use *U*=10,000 for the snRNA-seq data and *U*=80,000 for the 68K PBMC single-cell data, although this can be adjusted according to the dataset. We use 10,000 because this is a typical maximum number of cells that can be captured with current microfluidic devices^16^. The labels of test set droplets are allowed to change during the EM. We define the debris set as all the other droplets below the test set. We fix the assignments *z* of these low-count droplets to debris during the EM. Finally, the cluster set is used to estimate the number of cell types and assign droplets to those cell types for initialization. We define the cluster set as the top *C* droplets ordered by total number of genes detected. The number of *C* droplets should be selected so that the majority of them are expected to contain nuclei or cells. Here we use *C*=1,000 for the snRNA-seq data and *C*=30,000 for the 68K PBMC single-cell data. To remove lowly expressed genes, we calculate counts per million (CPM) for each gene in the test set and the debris set. Then, we only use expressed genes, which have CPM ≥ 10 in both the test set and debris set.

The number of mixtures and their parameter initializations are determined by running an initial clustering of the high-count droplets. A proper initialization is important because a multinomial mixture model is sensitive to different initializations^29^. To initialize the parameters for EM, we cluster the *C* droplets in the cluster set using the graph-based Louvain clustering algorithm^30^. First, we extract the top *V*=2,000 variable genes^19^. To do so, we take the filtered expressed genes and calculate the mean and variance of the raw gene counts within the cluster set. Then, we add a constant of 1 and log_10_ transform the means and variances. We then fit a loess regression line between the log transformed mean and variance values with a span=0.3. Finally, we take the top *V*=2,000 genes ranked by residual, which is calculated by subtracting the fitted variance from the observed variance. Then, we normalize the *V* x *C* raw count matrix to take into account total read depth and variance. The total number of UMI counts of the variable genes is calculated and then the droplet counts are divided by it so that the droplet counts sum to 1. The matrix is multiplied by the median number of counts across the *C* droplets. Finally, the adjusted counts are log transformed after adding a constant value of 1. This results in a normalized *V* x *C* count matrix. We construct a weighted k-nearest neighbors (kNN) graph using k=30 and weight=1/dist, where dist is the euclidean distance between a pair of droplets. The graph-based Louvain community detection algorithm^30^ is run on the kNN graph to identify clusters. Finally, we remove clusters containing less than 20 droplets. This provides the number of *k* cell types to use and the initialized droplet assignments. We use the mean gene counts of the droplets in cluster *k* to initialize *m*_*k*_ for EM. Finally, *π_k_* is initialized by dividing the total number of droplets in group *k* by the total number of droplets that have an assignment during the initialization.

After the initialization, we use semi-supervised EM to estimate the parameters *m*_*k*_ and *π_k_*. During the M step, we maximize the expected complete data log-likelihood with respect to the parameters. For *m*_*k*_, we add a pseudocount of 10^−4^ to avoid collapsing the likelihood to 0. During the E step, we calculate the probability of a droplet assigned to group *k*, followed by the expected value of *z* as the fraction of these probabilities. These two steps iterate during EM, and the algorithm converges when the change in likelihood is below ε, which we set to 10^−8^. We then assign droplets to debris if the probability of belonging to the debris group is greater than 0.95.

### Identifying cell types after filtering droplets

For all experiments, we ran a standardized clustering pipeline using Seurat v3.0.0^19^. After applying filtering, we only kept droplets with at least 200 genes detected^4^ to ensure that each droplet had enough information for clustering. The count data were log-normalized using the NormalizeData function in Seurat, using a scaling factor equal to the median of total counts across droplets. For the six adipose tissue samples, we used a scaling factor equal to 1,000 to ensure that all samples were normalized equally. Additionally, we merged the normalized data of the six adipose tissue samples without batch correction, as we saw high overlap of clusters among the six samples (data not shown). The top 2,000 variable genes were then calculated using the FindVariableFeatures function.

Normalized read counts for each gene were scaled to mean 0 and variance 1. We calculated the first 30 PCs to use as input for clustering. We then ran the Seurat functions FindNeighbors and FindClusters with 30 PCs. In the FindClusters function, we used the default parameters with standard Louvain clustering and a default clustering resolution of 0.8, unless otherwise stated. For visualization, we ran UMAP^22^ on the 30 PCs and set the spread parameter to 5. To identify marker genes for each cluster, we ran a Wilcoxon rank-sum test using the function FindAllMarkers with default parameters and only.pos=TRUE. We corrected for multiple testing using a false discovery rate (FDR) threshold of 0.05.

## Acknowledgements

We thank the Finnish individuals who participated in the adipose tissue study, as well as Jaakko Kaprio and Aila Rissanen for their contributions. We also thank the Finnish Institute for Molecular Medicine Single Cell Analytics Core for performing library preparation and sequencing for the adipose tissue samples. This study was funded by the National Institutes of Health (NIH) grants HL-095056, HL-28481, and U01 DK105561. M.A. was supported by the HHMI Gilliam Fellowship and the NIH T32HG002536. JRP, JNB and PP were supported by an NIH DK41301 grant. E.H., E.R., and B.J. were supported by the National Science Foundation grant 1705197. B.J. was supported by the National Science Foundation Graduate Research Fellowship Program under Grant No. DGE-1650604. K.M.G. was supported by the NIH F31HL142180. Z.M was supported by the AHA grant 19PRE34430112. K.H.P. was supported by the Academy of Finland (272376, 266286, 314383, 315035), Finnish Medical Foundation, Finnish Diabetes Research Foundation, Novo Nordisk Foundation, Gyllenberg Foundation, Sigrid Juselius Foundation, Helsinki University Hospital Research Funds, Government Research Funds and University of Helsinki.

## Data Availability

We will upload the human single nucleus RNA-seq data to GEO (accession number XX) upon acceptance of the manuscript.

## Code Availability

The DIEM program is freely available for use at https://github.com/marcalva/diem.

## Contributions

M.A., E.R., E.H., and P.P. conceived the study and designed the analysis. M.A., K.M.G., and J.B. designed, performed, and interpreted nuclei isolation experiments. M.A. and K.M.G. designed and performed snRNA-seq experiments and collected data. M.A., E.R. B.J., Z.M., K.M.G., and J.B. analyzed and interpreted snRNA-seq data. M.A., E.R., E.H. and P.P designed the DIEM method. J.R.P., C.J.Y., K.H.P. and P.P. designed and supervised nuclei isolation and snRNA-seq experiments. M.A., E.R., B.J., Z.M., E.H., and P.P. wrote the manuscript.

---

**Supplementary Figure 1. Extranuclear RNA can contaminate droplets with high UMI counts.**

Scatterplots showing the relationship between the percent of UMIs mapping to the mitochondrial genome (MT%) (y-axis) and the total UMI count in a droplet (x-axis) for the differentiating preadipocytes (DiffPA), mouse brain, and six human frozen adipose tissue (AT) snRNA-seq datasets. The dotted red line indicates the quantile-based threshold, calculated as the counts of the 99th percentile of expected nuclei (3,000) divided by 10.

**Supplementary Figure 2. Preservation of differential RNA profiles between nuclear-enriched and background-enriched droplets.**

Correlation plots of log fold changes across different snRNA-seq experiments. For each of the 8 experiments (differentiating preadipocytes (DiffPA), mouse brain, and six human frozen adipose tissue (AT) snRNA-seq samples), the log2 fold change of the counts per million (CPM) for each gene is calculated between the nuclear-enriched and background-enriched droplets. Nuclear-enriched and background-enriched droplets are those with UMI counts greater than or equal to, and less than 100 UMI counts, respectively.

**Supplementary Figure 3. DIEM preferentially removes contaminated droplets from frozen human adipose tissue.**

**a,b,c**, Clustering of droplets removed by **(a)** DIEM, **(b)** EmptyDrops^12^, and **(c)** quantile-based filtering methods. The left side shows a UMAP^22^ visualization of the removed droplets colored by cluster. The bar plots on the right show the percent of UMIs mapping to the mitochondrial genome (MT%) on the y-axis for droplets in each cluster (on the x-axis). The left bar plot shows clusters derived from removed droplets, while the right bar plot is derived from droplets passing the method’s filtering criteria.

**Supplementary Figure 4. Accurate modeling of major cell types used by the multinomial mixture model in DIEM.**

**a-h**, The left side shows scatter plots of a Principal Components Analysis (PCA) depicting the implicit cluster assignments by DIEM. The right side shows the overlap between the DIEM-assigned clusters and the Seurat-assigned clusters. The results are shows for the **(a)** differentiating preadipocytes (DiffPA), **(b)** the mouse brain, and **(c-h)** the six human frozen adipose tissue (AT) data sets. The scatter plots show PCs 1-2, 3-4, and 5-6, colored by DIEM-assigned cluster. The heatmaps on the right show the percent of droplets in the Seurat^19^ cluster (top) that belong to the DIEM cluster (left).

**Supplementary Figure 5. DIEM preserves the cell-types identified by established methods.**

**a**, Boxplots showing the percent of UMIs mapping to *MALAT1* (MALAT1%) per droplet in the differentiating preadipocytes (DiffPA). MALAT1% of clusters are compared across the quantile-based, EmptyDrops^12^, and DIEM filtering methods. **b**, Boxplots showing the total number of UMIs per droplet in the differentiating preadipocytes (DiffPA). Clusters are compared across the three filtering methods.

**Supplementary Figure 6. DIEM preserves the cell-types identified by established methods.**

**a,b**, The correspondence between quantile/EmptyDrops^12^ clusters and DIEM clusters are shown for the **(a)** mouse brain and **(b)** adipose tissue (AT) data sets. Quantile and EmptyDrops clusters are identified using a resolution of 0.8, while DIEM clusters are identified with a higher resolution of 2.0 with Seurat^19^. Each circle is a cluster identified after filtering with the method specified at the top. The thickness of a connecting line between a quantile/EmptyDrops cluster and a DIEM cluster represents the percent of droplets in the quantile/EmptyDrops cluster that are contained in the connecting DIEM cluster. No lines emerging from a quantile/EmptyDrops cluster means that droplets from that cluster are removed by DIEM. The color of a cluster circle indicates the average percent of mitochondrial UMIs per cluster (MT%).

**Supplementary Figure 7. DIEM reduces the total number of MT marker genes in tissue-based snRNA-seq.**

Bar plots showing the total number of mitochondrial marker genes across all clusters for the differentiating preadipocytes (DiffPA), mouse brain, and human frozen adipose tissue (AT) data sets. The total number of MT markers is defined as the sum of the number of statistically significant MT markers in a cluster across all clusters.

