## SupplementaryFigures for "Enhancing droplet-based single-nucleus RNA-seq resolution using the semi-supervised machine learning classifier DIEM"

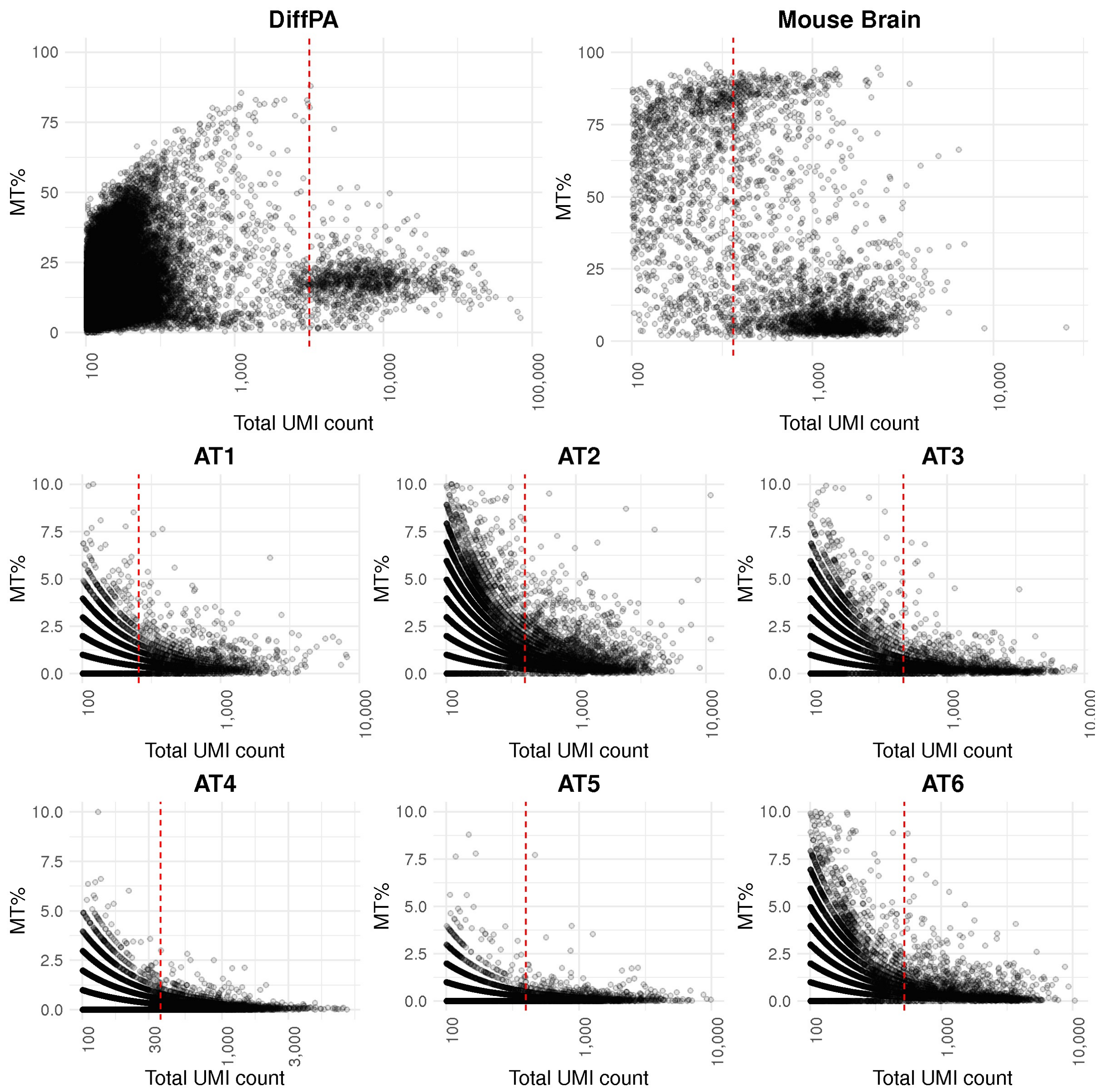


**Supplementary Figure 1. Extranuclear RNA can contaminate droplets with high UMI counts.**

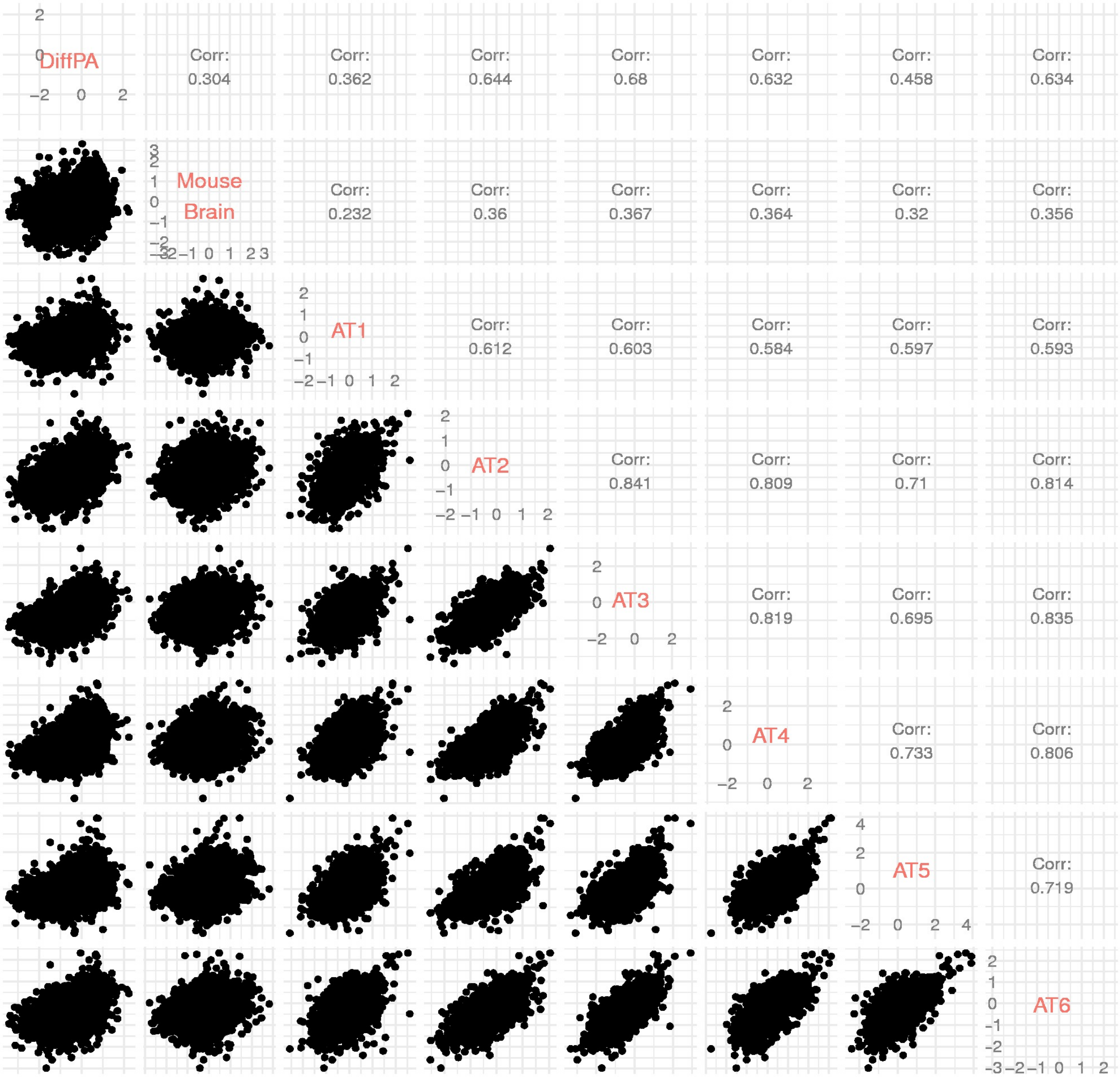


**Supplementary Figure 2. Preservation of differential RNA profiles between nuclear-enriched and background-enriched droplets.**

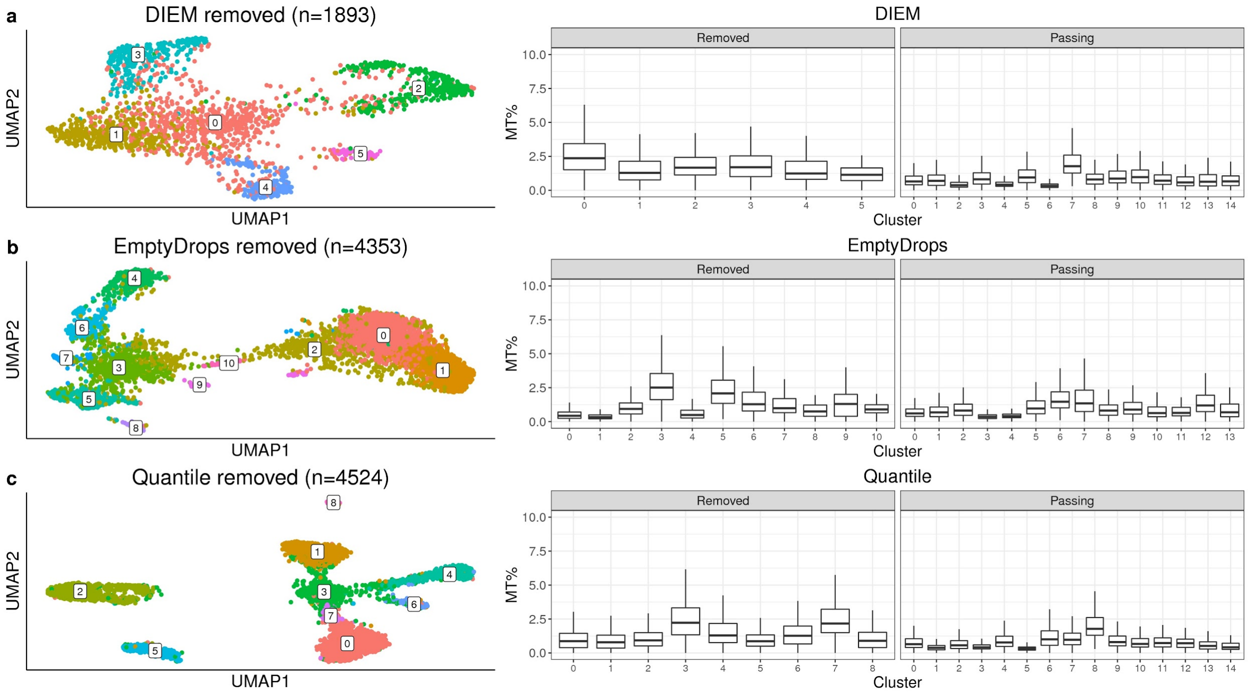


**Supplementary Figure 3. DIEM preferentially removes contaminated droplets from frozen human adipose tissue.**

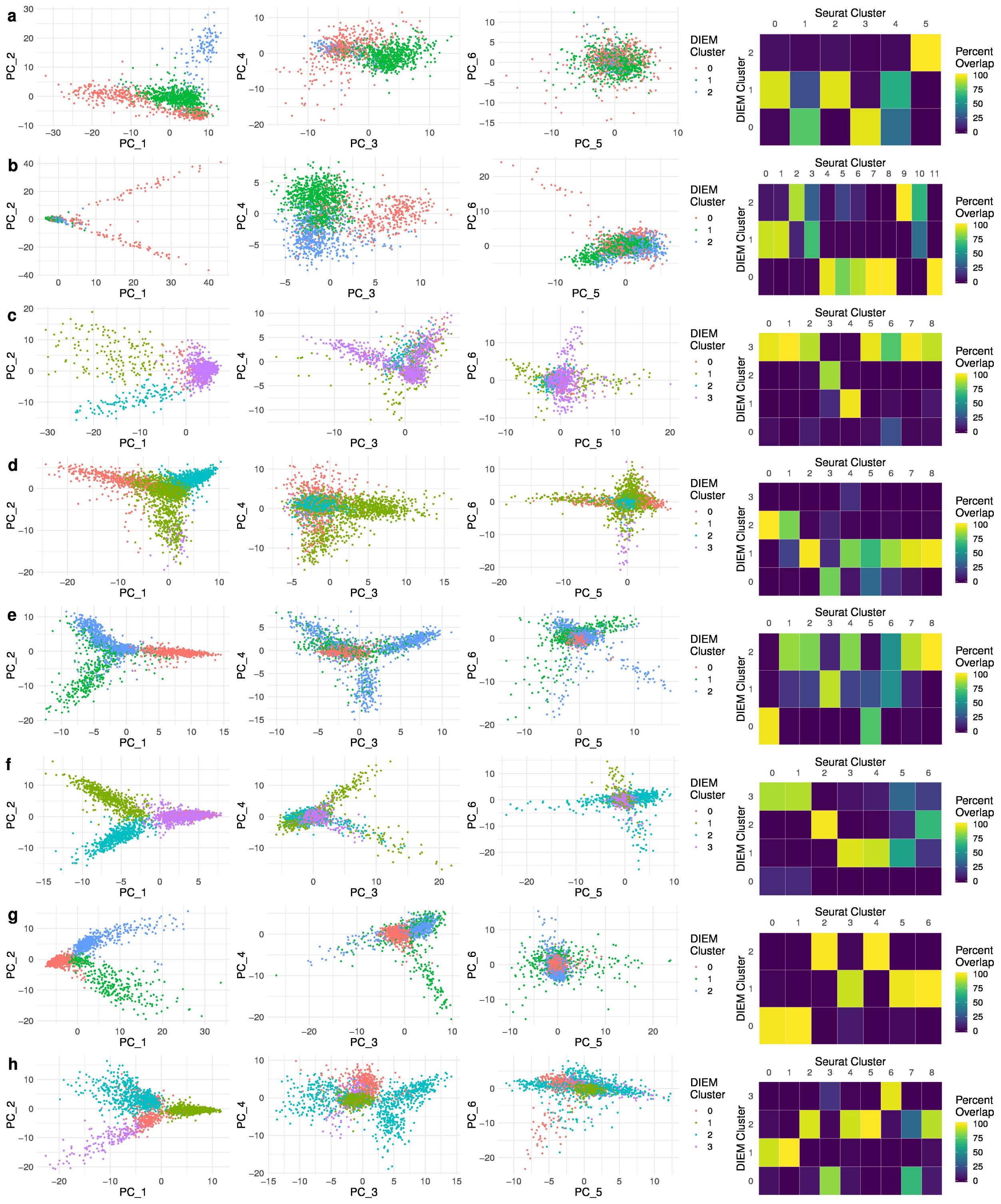


**Supplementary Figure 4. Accurate modeling of major cell types used by the multinomial mixture model in DIEM.**

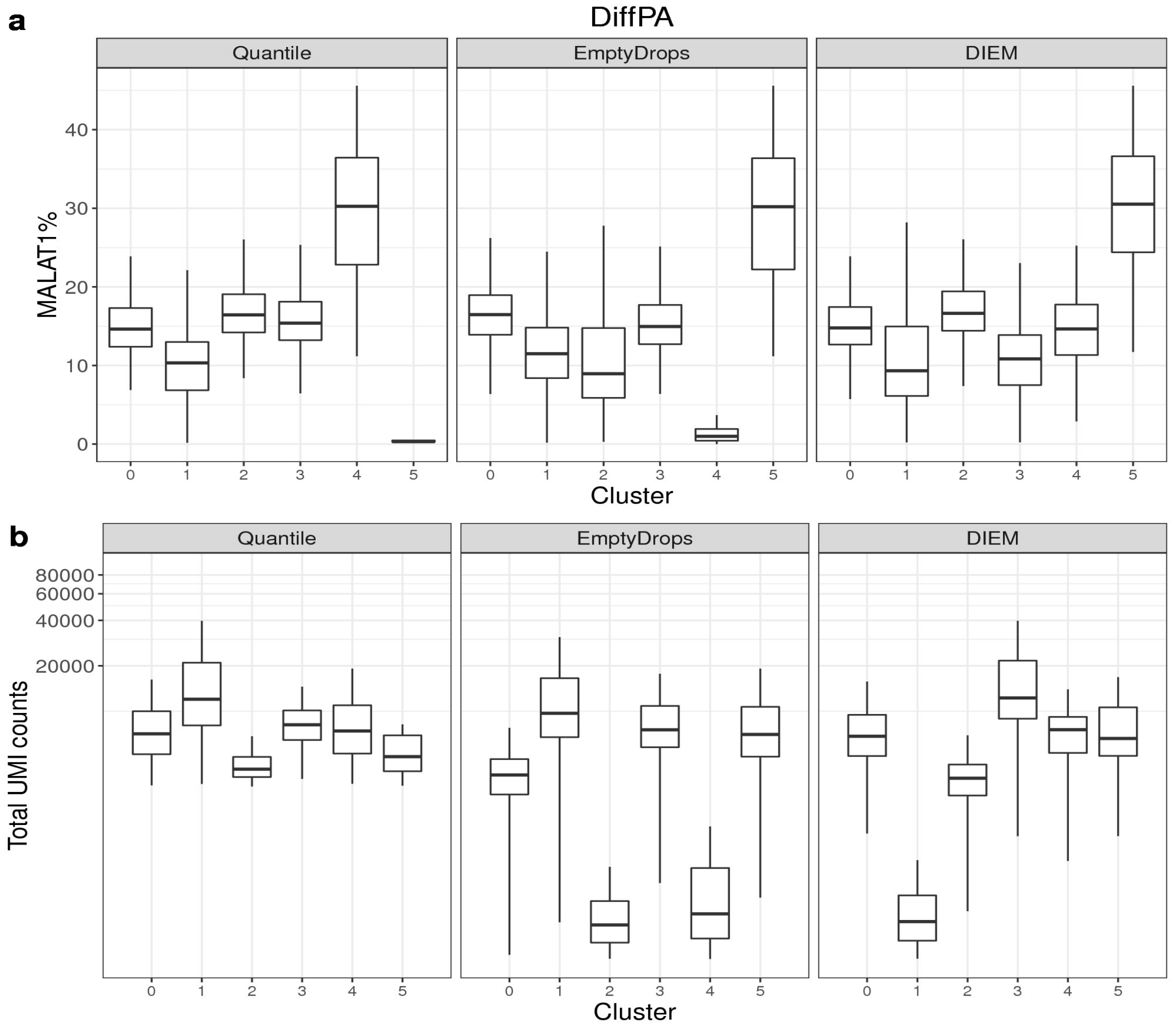


**Supplementary Figure 5. DIEM preserves the cell-types identified by established methods.**

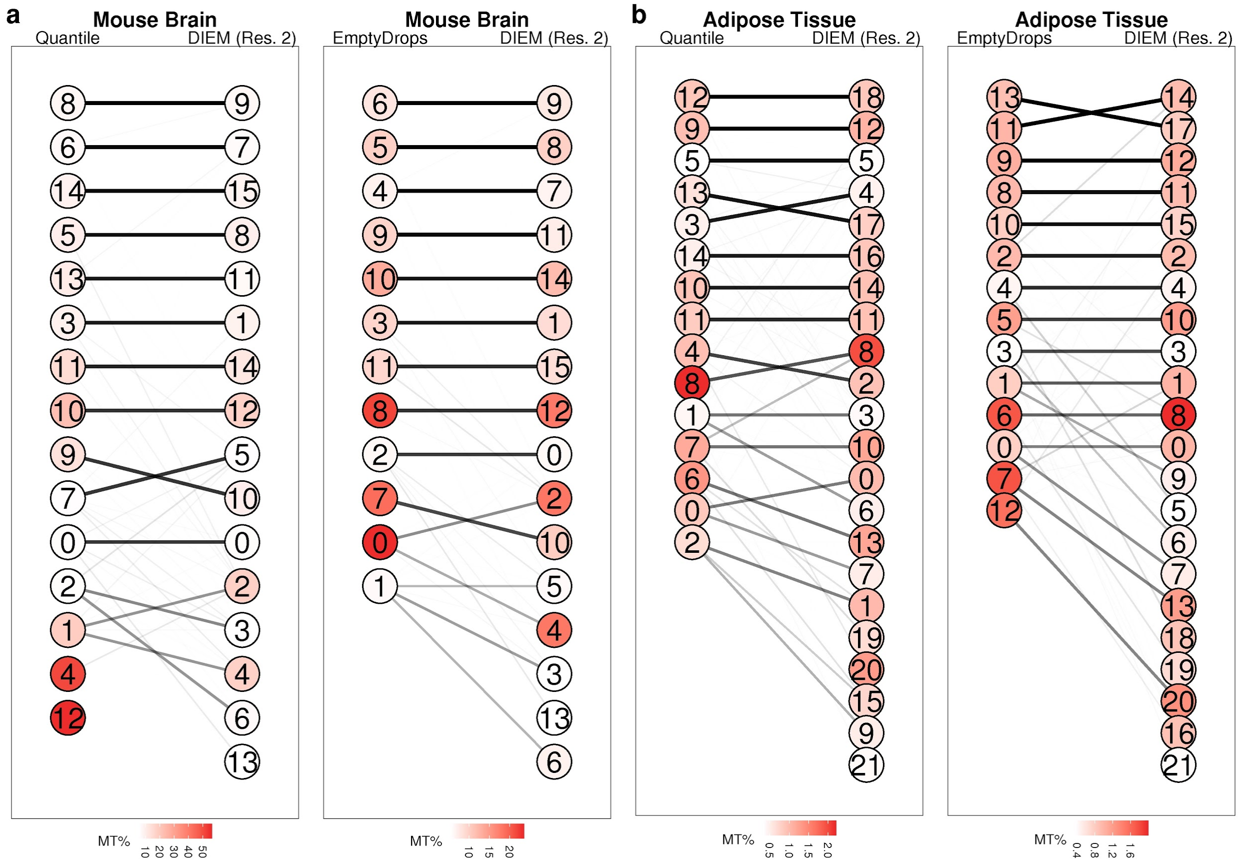


**Supplementary Figure 6. DIEM preserves the cell-types identified by established methods.**

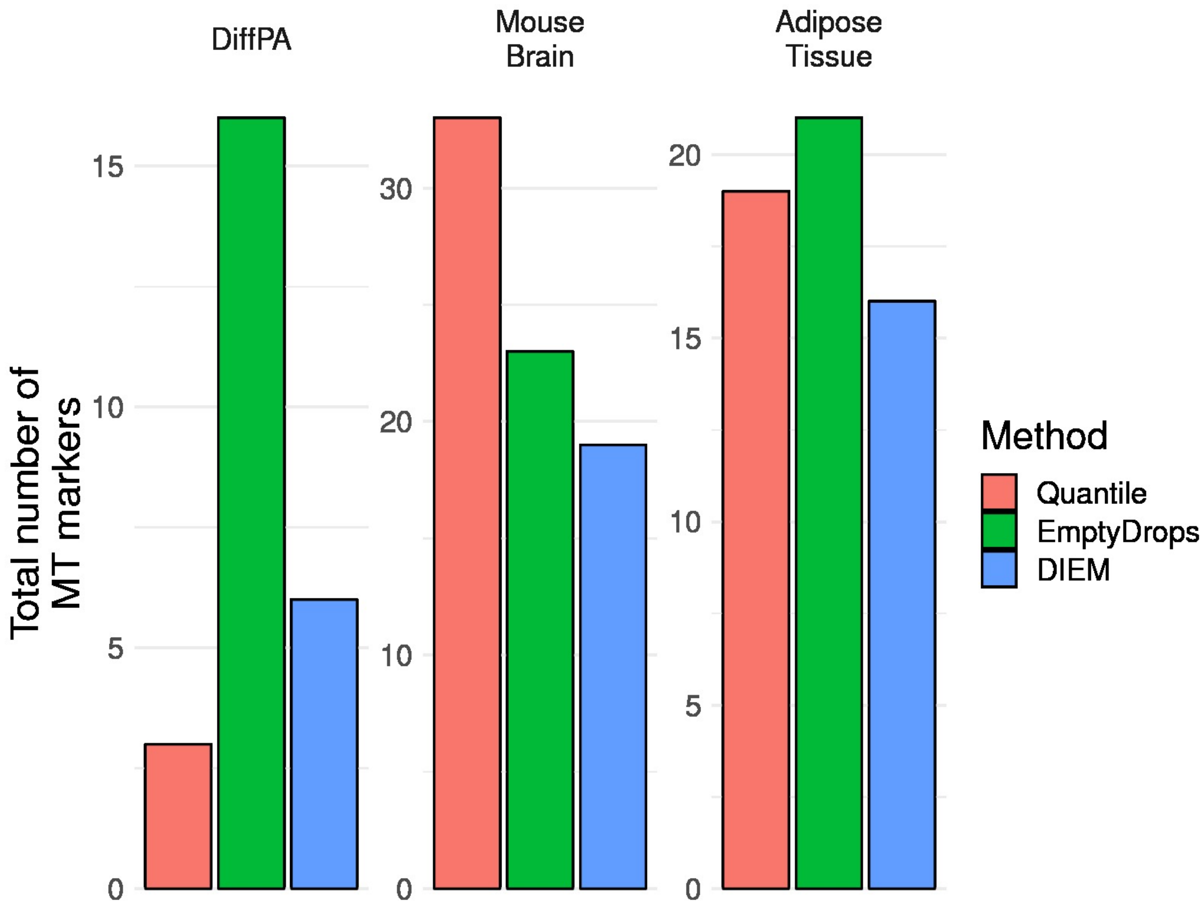


**Supplementary Figure 7. DIEM reduces the total number of MT marker genes in tissue-based snRNA-seq.**
